# Increased autonomous bioluminescence emission from mammalian cells by enhanced cofactor synthesis

**DOI:** 10.1101/2024.08.13.607744

**Authors:** Theresa Brinker, Carola Gregor

## Abstract

The bacterial bioluminescence system has been successfully implemented in mammalian cell lines, enabling substrate-free luminescence imaging of living cells. One of the major limitations of the system is its comparatively low brightness. To improve light emission, we aimed at increasing the cellular production of FMNH_2_ and NADPH which serve as cosubstrates in the bacterial bioluminescence reaction. We coexpressed different proteins involved in the synthesis of these two cofactors together with the proteins of the bacterial bioluminescence system in different mammalian cell lines. Combined expression of a riboflavin kinase (RFK) and a constitutively active Akt2 variant (Akt2CA) that participate in the cellular production of FMN and NADP^+^, respectively, increased bioluminescence emission up to 2.4–fold. The improved brightness allows autonomous bioluminescence imaging of mammalian cells at a higher signal-to-noise ratio and enhanced spatiotemporal resolution.

## 1. Introduction

Optical imaging of living cells, tissues and organisms can be implemented by detecting photons generated through bioluminescence. In this process, light is produced by the sample itself through a biochemical reaction. Bioluminescence imaging has the advantage that no light source is required and thus imaging can be performed at exceedingly low light levels. Hence, light-induced effects such as phototoxicity, photobleaching or the pertubation of light-sensitive cells which can be problematic in fluorescence measurements do not occur. Additionally, bioluminescence offers the advantage of a very low background signal due to the absence of autofluorescence.

The generation of bioluminescence light is based on the oxidation of a small-molecular substrate, the luciferin. This reaction is catalyzed by an enzyme, the luciferase. Luciferases from different species found in nature (e.g., firefly luciferase (FLuc) or *Renilla* luciferase (RLuc)) and engineered luciferases (e.g., NanoLuc (NLuc) [1]) can be heterologously expressed in different cell types, particularly in mammalian cells. Addition of the appropriate luciferin results in cellular light production that allows image acquisition without any additional interventions in the sample.

External luciferin supply is associated with several problems such as limited cell permeability and biodistribution of the luciferin [2,3], elevated background signal due to its autooxidation in solution [4], decay of the signal during long-term measurements [5,6] and high costs. Therefore, biochemical pathways for cellular synthesis and recycling of the luciferin have been explored. Coexpression of their enzymes together with the luciferase enables autonomous bioluminescence imaging of the target cells without the need for an external substrate. Up to now, the enzymes for luciferin production have only been identified in their entirety for two bioluminescence systems, which are those from bioluminescent bacteria [7–11] (Figure 1A) and fungi [12]. Of these two, only the bacterial system is fully functional in mammalian cells [13–15] which are of greatest interest for cell biological and medical issues. Applications of this autonomous bioluminescence system are mainly limited by its lower brightness compared to other luciferases that require an external luciferin.

**Figure 1.**
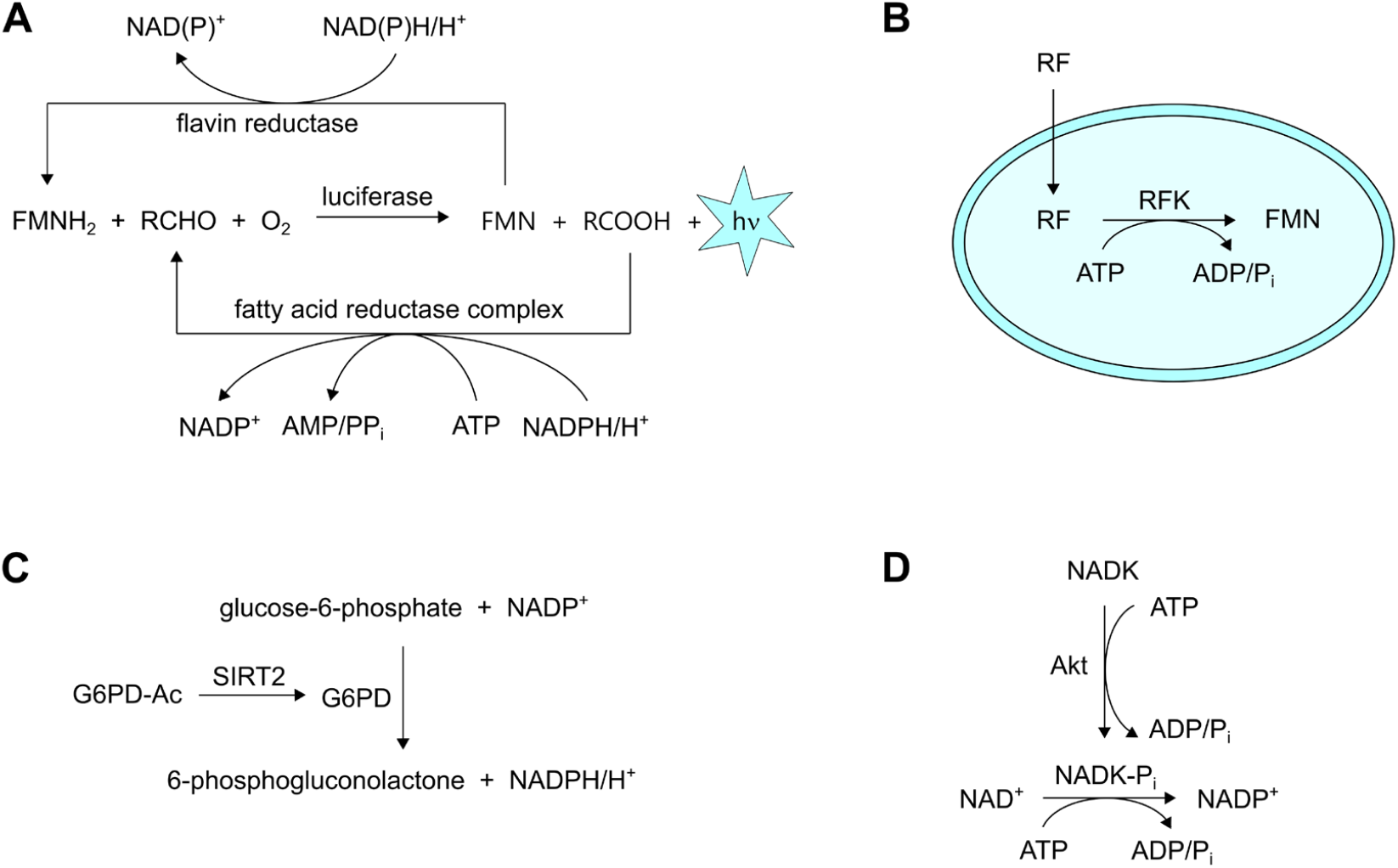
Reactions involved in light generation by the bacterial bioluminescence system in mammalian cells. (**A**) Bacterial luciferase catalyzes the oxidation of FMNH_2_ and a fatty aldehyde (RCHO) to FMN and a fatty acid (RCOOH) with concomitant photon emission. The two products are regenerated by a flavin reductase and the fatty acid reductase complex, respectively. (**B**) Riboflavin (RF) is absorbed by the cell and converted into FMN by riboflavin kinase (RFK). (**C**) Glucose-6-phosphate dehydrogenase (G6PD) catalyzes the reduction of NADP^+^ to NADPH. The enzyme is regulated by sirtuin 2 (SIRT2) that deacetylates and thereby activates G6PD. (**D**) NADP^+^ is produced from NAD^+^ by NAD kinase (NADK). NADK is activated by phosphorylation by protein kinase B (Akt).

The light levels produced by the bacterial bioluminescence system do not only depend on the activity of the luciferase, but also on the availability of its substrates. We therefore set out to improve light generation in mammalian cells by increasing the cellular synthesis of two cosubstrates. As the first target compound, we chose flavin mononucleotide (FMN) that is directly involved in the bacterial bioluminescence reaction. Our second target was NADPH which serves as the reductant for the regeneration of the two substrates of the bioluminescence reaction, reduced FMN (FMNH_2_) and a long-chain fatty aldehyde (Figure 1A). By overexpression of two enzymes involved in the respective synthetic pathways (Figure 1B– D), autonomous bioluminescence from human cell lines was increased up to 2.4–fold. The enhanced brightness enables single-cell imaging by bioluminescence microscopy with improved spatiotemporal resolution and signal-to-noise ratio (SNR).

## 2. Materials and Methods

### 2.1 Generation of plasmids

To generate plasmids for expression in mammalian cells, genes were PCR-amplified with the primers listed in Table S1 using the following templates: RFK: pDONR223-RFK (Addgene plasmid #23698) [16], G6PD: G6PD/pRK5 (Addgene plasmid #41521) [17], SIRT2: SIRT2 Flag (Addgene plasmid #13813) [18], NADK: pDONR223-NADK (Addgene plasmid #23747) [16], Akt1 and Akt1CA: pDONR223-AKT1 (Addgene plasmid #23752) [16], Akt2 and Akt2CA: pLenti-FLAG-Akt2-CA (Addgene plasmid #64050) [19]. The PCR products were cloned into pcDNA3.1(+) between the NheI and XhoI restriction sites (NheI and BamHI sites for NADK). For simultaneous expression of RFK and Akt2CA from a single vector, plasmids containing Akt2CA-P2A-RFK and RFK-P2A-Akt2CA were produced. Both genes were PCR-amplified separately using the primers listed in Table S1. The first insert was digested with NheI and BamHI and the second insert with BamHI and XhoI. Both fragments were ligated simultaneously into pcDNA3.1(+) digested with NheI and XhoI.

For measurements of the NADPH concentration in mammalian cells, the plasmid iNap1 pcDNA3.1 Hygro(+) [20] was used. For bacterial expression, His-iNap1 was PCR-amplified from iNap1 pcDNA3.1 Hygro(+) using the primers listed in Table S1 and inserted into the expression vector pQE(-) [21] between the BamHI and SalI restriction sites.

### 2.2 Cell culture and transfection

LiveLight HEK293 cells were obtained from 490 BioTech (Knoxville, TN, USA). HEK293 cells were obtained from DSMZ (German Collection of Microorganisms and Cell Cultures GmbH, Braunschweig, Germany, cat. no. ACC 305). HeLa cells were obtained from LGC Standards (Wesel, Germany, cat. no. ATCC-CCL-2). All cells were cultured at 37 °C in 5% CO__2__ in DMEM with 4.5 g/l glucose (Thermo Fisher Scientific, Waltham, MA, USA) supplemented with 10% FBS (Merck KGaA, Darmstadt, Germany), 1 mM sodium pyruvate (Thermo Fisher Scientific), 100 units/ml penicillin and 100 μg/ml streptomycin (Merck).

Cells grown in 24-well plates with 0.5 ml of cell culture medium were transfected with a total amount of 0.5 µg DNA per well using 1 µl jetPRIME transfection reagent (Polyplus-transfection, Illkirch, France) according to the instructions of the manufacturer. For expression of Lux proteins, a mixture of plasmids (hereafter referred to as lux plasmids) containing the individual codon-optimized genes at a *luxA*:*luxB*:*luxC*:*luxD*:*luxE*:*frp* ratio of 1:1:3:3:3:1 was used as described previously [15]. Additional plasmids were transfected simultaneously in the indicated quantities.

For microscopy, cells were grown on 18 mm round coverslips (VWR, Radnor, PA, USA) in 12-well plates with 1 ml of cell culture medium per well and transfected with a total amount of 1 µg DNA and 2 µl jetPRIME as described above. The medium was exchanged against fresh cell culture medium 4 hours post-transfection.

### 2.3 Bioluminescence imaging

For comparison of the bioluminescence emission of cells transfected with different plasmids, cells were imaged in 24-well plates with an Amersham Imager 680 RGB (Cytiva, Marlborough, MA, USA) 24 hours post-transfection.

Bioluminescence microscopy was conducted with a custom-built setup as described previously [21,15]. A PlanApo N 60x/1.42 oil objective lens (Olympus, Tokio, Japan) was used, resulting in an effective pixel size on the camera of 360 nm. Cells were kept at 37 °C in 5% CO_2_ during imaging.

### 2.4 Fluorescence measurements of iNap1

Fluorescence measurements of iNap1 were performed with the same microscope as for bioluminescence imaging using lasers at 405 and 491 nm for excitation. Measurements of mammalian cells were conducted at 37 °C in 5% CO_2_.

A calibration curve was recorded using purified His-tagged iNap1 protein. The protein was expressed from the His-iNap1 pQE(-) plasmid in *E. coli* Top10 cells. Cells were grown in LB medium to an OD_600_ of 0.6 and induced with 1 mM IPTG overnight at 30 °C. The following day, cells were harvested by centrifugation and resuspended in 4 ml B-PER Bacterial Protein Extraction Reagent (Thermo Fisher Scientific) per gram of cell pellet containing 0.1 mg/ml lysozyme (Merck), 10 U/ml DNaseI (Thermo Fisher Scientific), 200 µM PMSF (Thermo Fisher Scientific) and 1 µg/ml pepstatin A (Merck). After incubation at room temperature for 20 min, 2 ml binding buffer (20 mM sodium phosphate, 500 mM NaCl, 20 mM imidazole, pH 7.4) were added per ml of B-PER. The solution was centrifuged for 20 min at 2000 g to remove cell debris. The supernatant was applied to a HisPur Ni-NTA spin column (Thermo Fisher Scientific) according to the manufacturer’s directions and His-iNap1 was eluted with 500 µl elution buffer (20 mM sodium phosphate, 500 mM NaCl, 500 mM imidazole, pH 7.4). The buffer was exchanged to 100 mM HEPES, 100 mM KCl, pH 7.3 by ultrafiltration using Vivaspin 500 spin concentrators (10 kDa MWCO, Merck). Fluorescence measurements were performed at 37 °C in the same buffer supplemented with different concentrations of NADPH (Carl Roth, Karlsruhe, Germany) on the microscope using the same settings as for cell imaging.

### 2.5 MTT assay

Cells were seeded into 24-well plates and transfected with the indicated plasmids as described above. 24 hours post-transfection, 50 μL of MTT (Merck) in PBS (5 mg/ml) were added. The cells were incubated at 37 °C in 5% CO_2_ for 60 min before addition of 0.5 ml SDS (Merck) solution (10%). The cells were resuspended by pipetting and absorption at 570 nm was measured with a NanoPhotometer N60 UV/Vis (Implen, München, Germany).

## 3. Results

### 3.1. Influence of individual proteins on bioluminescence intensity

In the first step, we aimed to increase the cellular production of FMNH_2_ which is a direct substrate of the bacterial bioluminescence reaction. FMNH_2_ is produced from FMN by a flavin reductase (Figure 1A) that we coexpressed together with the other proteins of the bacterial bioluminescence system as described previously [15]. To enhance the concentration of the FMN precursor, we now additionally overexpressed human riboflavin kinase (RFK). This enzyme generates FMN by phosphorylation of riboflavin (Figure 1B) which is taken up into the cell by specific transporters [22–24]. Since animals are unable to produce riboflavin, no pathways for its synthesis exist in mammalian cells that could provide another possible target to increase FMN production. Coexpression of RFK together with the proteins of the bacterial bioluminescence system in different human cell lines resulted in increased bioluminescence emission (Figure 2). The largest improvement in bioluminescence intensity (more than 2–fold) was obtained in the autobioluminescent cell line LiveLight HEK293 (Figure 2A), whereas the brightness increased by ∼30 and 45% in unmodified HEK293 and HeLa cells, respectively (Figure 2B,C). We attribute the larger improvement in light emission from LiveLight HEK293 cells to the fact that a 5–fold larger amount of the RFK plasmid was applied since all genes of the bacterial bioluminescence system are constitutively expressed in this cell line and hence do not need to be cotransfected.

**Figure 2.**
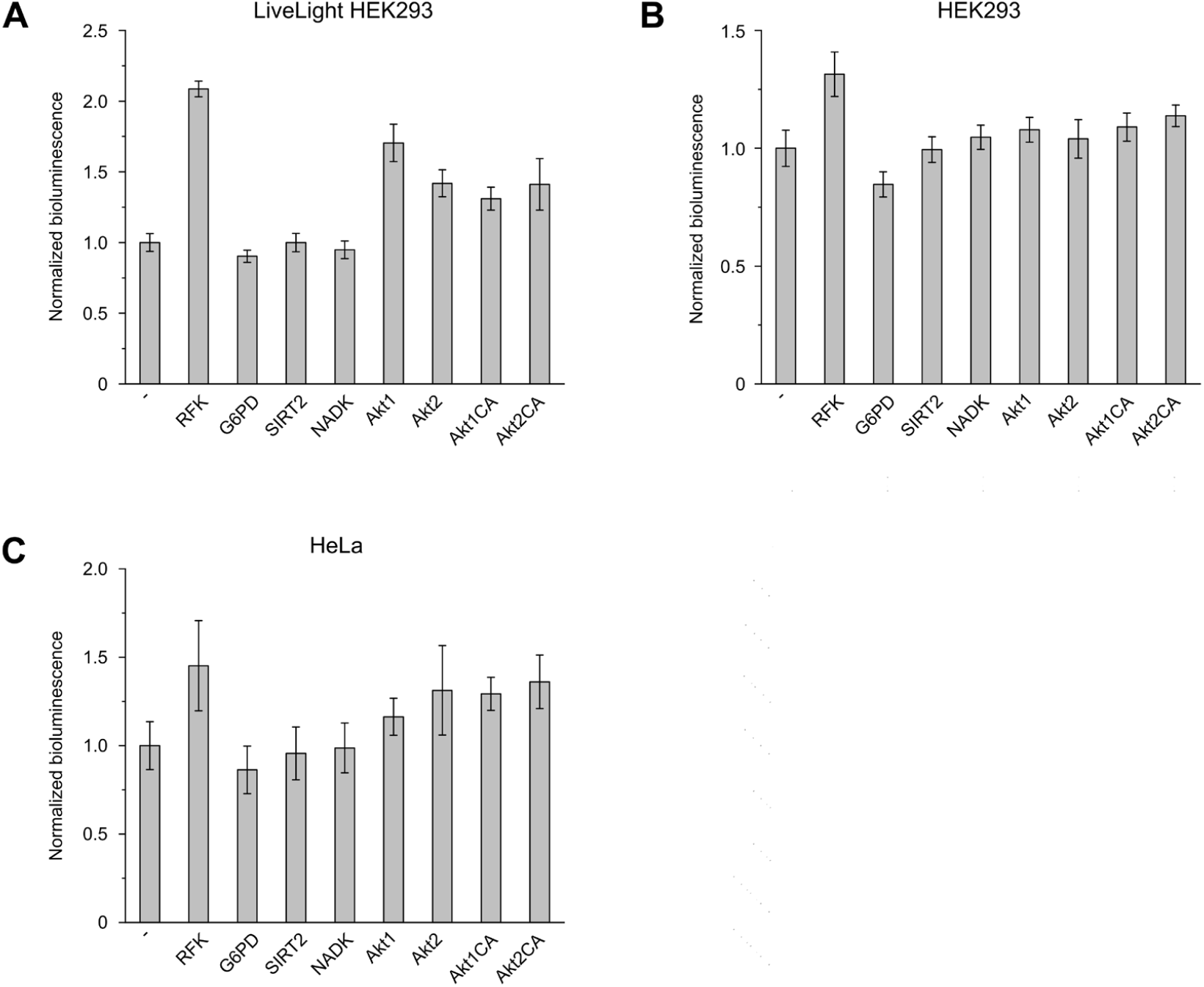
Effect of overexpression of different proteins on the bioluminescence emission in mammamlian cell lines: (**A**) LiveLight HEK293; (**B**) HEK293; (**C**) HeLa. Cells grown in 24-well plates were transfected with a mixture of 0.4 µg lux plasmids and 0.1 µg of the indicated genes (all constructs in pcDNA3.1(+)). The signal was normalized to the bioluminescence emission of cells transfected with 0.4 µg lux plasmids and 0.1 µg of the empty pcDNA3.1(+) vector (-). For LiveLight HEK293, 0.5 µg of the indicated gene in pcDNA3.1(+) was transfected. Error bars represent standard deviation from 5 wells.

In the second step, we investigated several enzymes involved in cellular NADPH synthesis (Figure 1C,D). We first chose glucose-6-phosphate dehydrogenase (G6PD) which is the rate-limiting enzyme of the pentose phosphate pathway and produces the majority of cytosolic NADPH by reduction of NADP^+^ [25]. Overexpression of G6PD did not increase bioluminescence emission in any of the cell lines tested (Figure 2). Likewise, no improvement in brightness was observed upon overexpression of sirtuin 2 (SIRT2) that regulates G6PD activity by deacetylation [26] (Figure 1C). This could be explained by the fact that cells maintain an NADPH:NADP^+^ ratio of around 100 [27,28], and hence a further increase of this ratio has only a minor influence on the NADPH concentration. Therefore, we next tested if higher bioluminescence emission can be achieved by increasing the total cellular concentration of NADPH and NADP^+^. Since NADPH is produced from NADP^+^, we aimed at enhancing NADP^+^ synthesis by increasing the reaction rate of NAD kinase (NADK) that catalyzes the conversion of NAD^+^ into NADP^+^ (Figure 1D). To this end, we overexpressed NADK which has been reported to increase cellular NADPH levels 4–5 fold [29]. Contrary to expectations, we observed no improvement of the bioluminescence signal upon NADK expression (Figure 2), possibly as a result of different cellular mechanisms that regulate the activity of this enzyme [30]. As an alternative strategy, we overexpressed protein kinase B (Akt) instead which activates NADK by phosphorylation [31] (Figure 1D). Expression of different Akt variants increased light emission in all tested cell lines (Figure 2). The largest effect in unmodified HEK293 and HeLa cells was observed for the constitutively active Akt2 variant Akt2CA [19] which we therefore selected for further measurements.

To verify the effect of Akt2CA expression on the cellular NADPH concentration, we compared cytosolic NADPH levels of cells with and without Akt2CA using the fluorescent sensor iNap1 [20]. iNap1 is a ratiometric sensor that increases its fluorescence excited at 420 nm upon binding of NADPH whereas the fluorescence excited at 485 nm decreases [20]. We determined the ratio of iNap1 fluorescence excited at 405 and 491 nm in Lux-expressing cells with and without Akt2CA (Figure 3). In all cell lines tested, expression of Akt2CA resulted in a significant increase of the fluorescence ratio, indicating an enhanced cellular NADPH concentration (Figure S1, Table S2) which confirms the anticipated effect of Akt2CA on NADPH synthesis.

**Figure 3.**
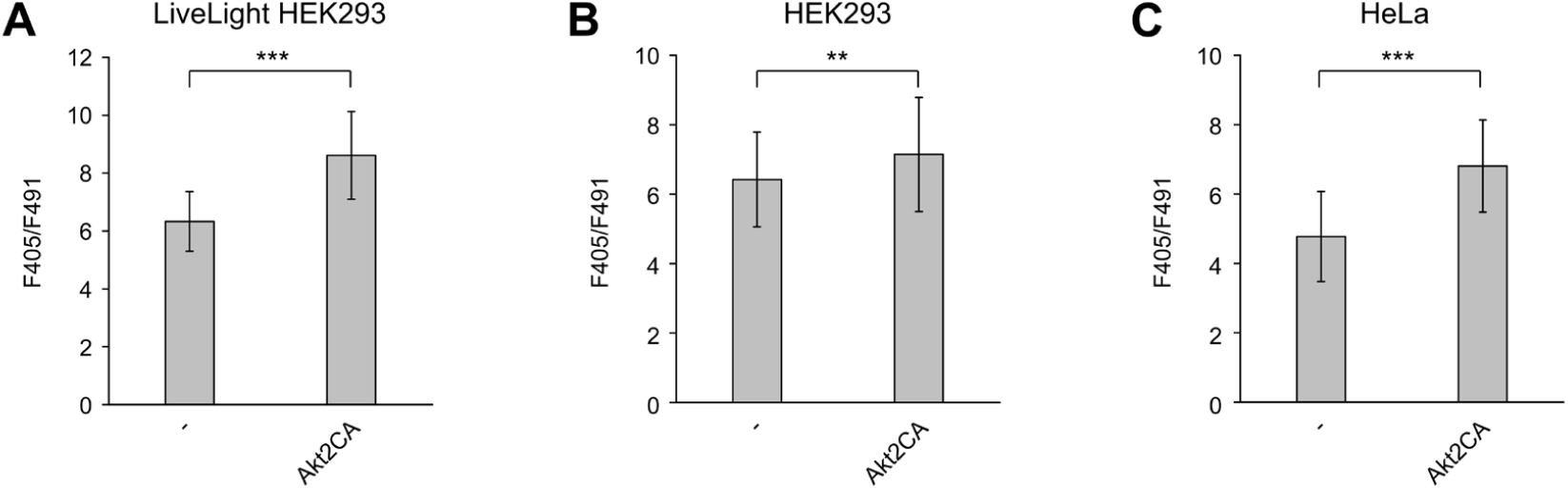
Fluorescence ratio of iNap1 in Lux-expressing cells with and without Akt2CA. (**A**) LiveLight HEK293; (**B**) HEK293; (**C**) HeLa cells grown on coverslips were transfected with a mixture of 0.6 µg lux plasmids, 0.2 µg iNap1 pcDNA3.1 Hygro(+) and 0.2 µg Akt2CA pcDNA3.1(+) or the empty pcDNA3.1(+) vector (-). For LiveLight HEK293, 0.2 µg iNap1 pcDNA3.1 Hygro(+) and 0.8 µg Akt2CA pcDNA3.1(+) or the empty pcDNA3.1(+) vector was transfected. iNap1 fluorescence excited at 405 (F405) and 491 nm (F491) was recorded with a custom-built microscope. Error bars represent standard deviation from at least 50 cells. ** and *** represent *P* values of <0.01 and <0.001, respectively, calculated by a 2-tailed Student’s *t* test.

### 3.2. Combined expression of RFK and Akt2CA

To combine the effects of RFK and Akt2CA on bioluminescence emission, we cotransfected both plasmids together with the lux plasmids. Since only a limited amount of DNA can be transfected, we reduced to half the quantity of each plasmid for their cotransfection to keep the total DNA amount constant. Combined expression of RFK and Akt2CA improved bioluminescence emission in HEK293 and HeLa cells compared to the individual plasmids transfected in double quantity whereas in LiveLight HEK293 cells the brightness with the RFK plasmid alone was slightly higher (Figure 4). Since the effects of RFK and Akt2CA on bioluminescence emission are expected to depend on their expression levels, we increased the transfected copy number of both genes by expressing them from a single plasmid. For this purpose, the RFK and Akt2CA genes were separated by a P2A sequence. Since the P2A peptide can affect protein function and hence also the brightness, we tested both orders of the two genes (Akt2CA-P2A-RFK and RFK-P2A-Akt2CA). In all cell lines, the highest brightness was obtained with Akt2CA-P2A-RFK, resulting in an up to 2.4–fold increase in bioluminescence emission (Figure 4).

**Figure 4.**
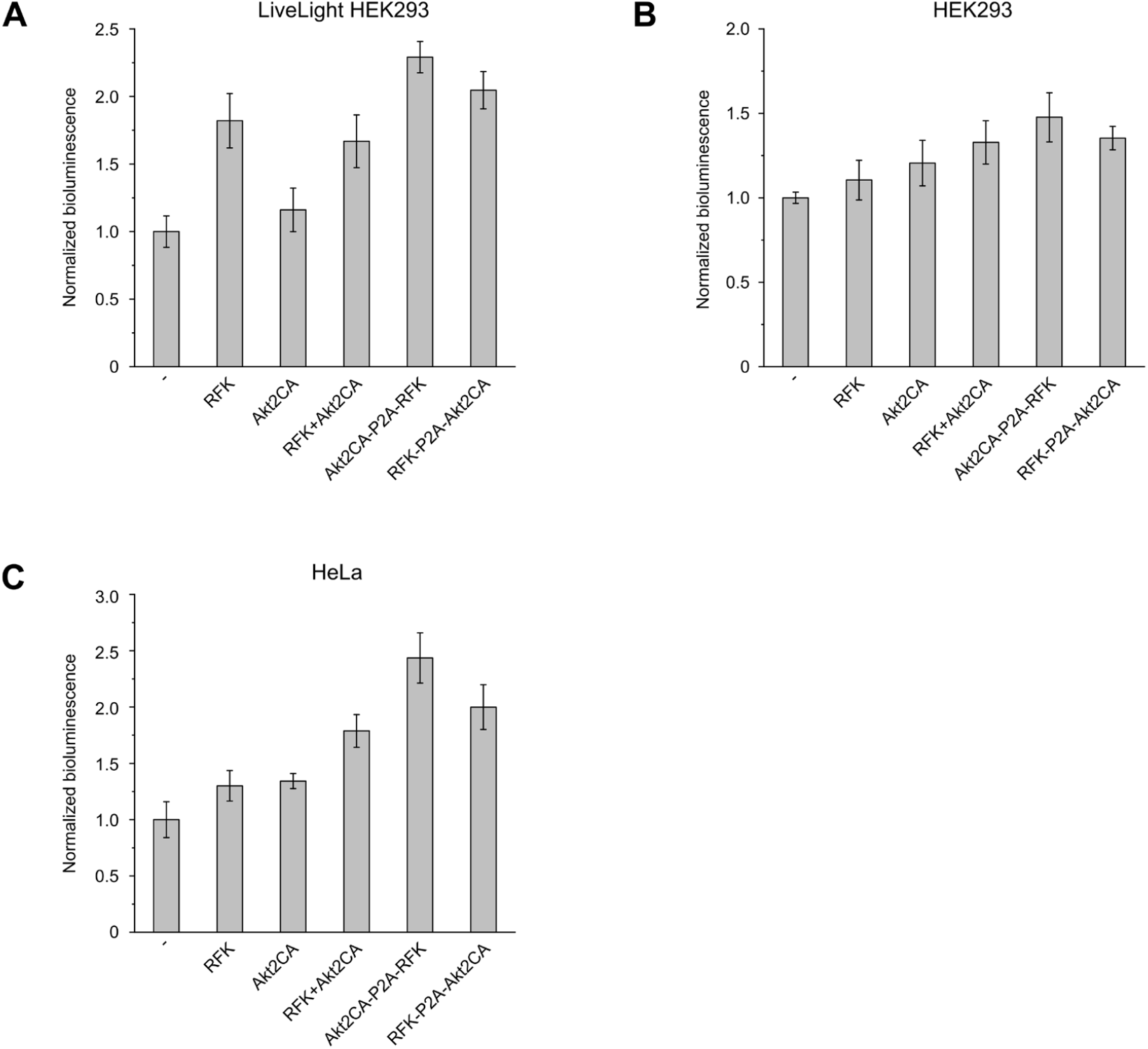
Effect of combined expression of RFK and Akt2CA on the bioluminescence emission in different mammalian cell lines: (**A**) LiveLight HEK293; (**B**) HEK293; (**C**) HeLa. Cells were grown in 24-well plates and transfected with a mixture of 0.4 µg lux plasmids and 0.1 µg of the indicated constructs (all in pcDNA3.1(+)). For RFK+Akt2CA, two separate plasmids containing RFK and Akt2CA were cotransfected (0.05 µg each). The signal was normalized to cells transfected with 0.4 µg lux plasmids and 0.1 µg of the empty pcDNA3.1(+) vector (-). For LiveLight HEK293, 0.5 µg of the indicated constructs was transfected. Error bars represent standard deviation from 5 wells.

We next investigated the optimal ratio of the lux and Akt2CA-P2A-RFK plasmids. Highest light levels were obtained with an Akt2CA-P2A-RFK percentage of 10 to 30% in both HEK293 and HeLa cells (Figure S2). We therefore used a mixture of 20% Akt2CA-P2A-RFK and 80% lux plasmids for the following experiments.

To assess possible effects of Akt2CA-P2A-RFK expression on cell viability, we performed MTT assays of cells transfected with Akt2CA-P2A-RFK or the empty vector (Figure 5). While reduced cell viability would be reflected by a decreased formazan formation and hence lower absorbance at 570 nm, we instead observed a slight increase in absorbance in all cell lines. This effect may result from the dependence of MTT reduction on NADPH-dependent oxidoreductases [32] and increased cellular NADPH production by Akt2CA. Overall, no negative effect of Akt2CA-P2A-RFK expression on cell viability or metabolic activity was detected.

**Figure 5.**
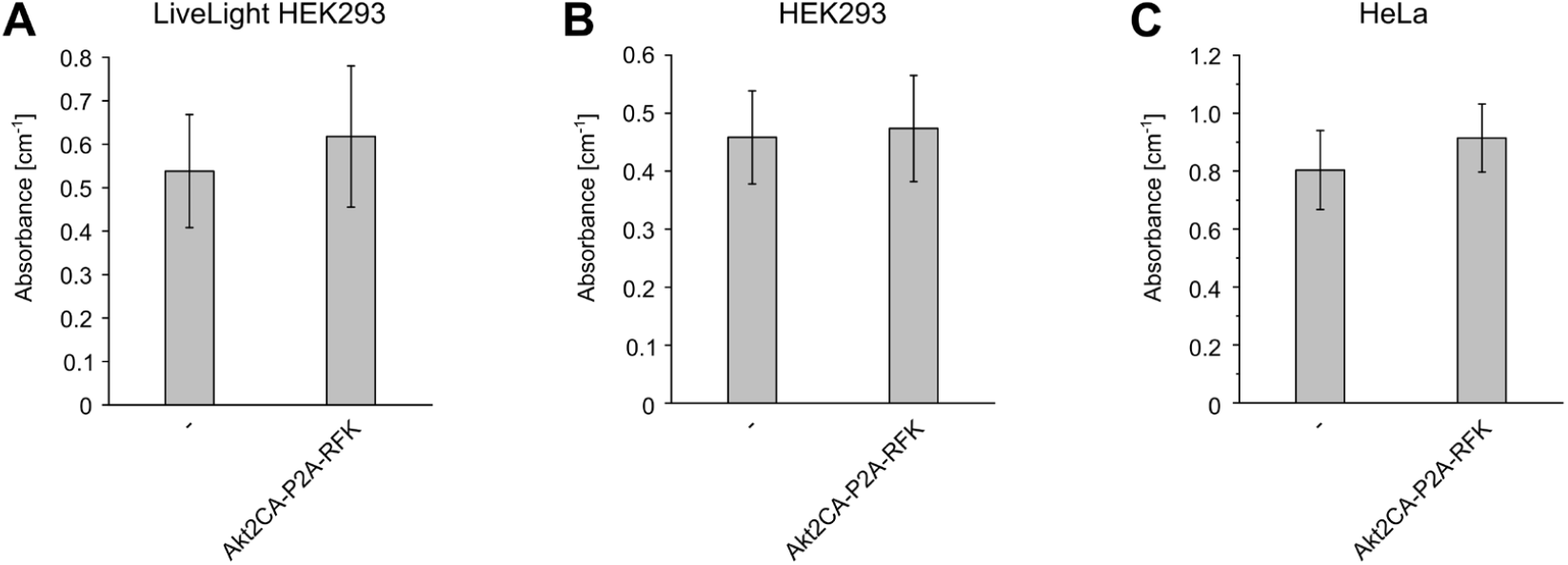
MTT assay of cells transfected with Akt2CA-P2A-RFK. (**A**) LiveLight HEK293; (**B**) HEK293; (**C**) HeLa cells grown in 24-well plates were transfected with a mixture of 0.1 µg Akt2CA-P2A-RFK and 0.4 µg of the empty vector or 0.5 µg of the empty vector (-) (all constructs in pcDNA3.1(+)). For LiveLight HEK293, 0.5 µg of the empty vector or the Akt2CA-P2A-RFK plasmid was transfected. Absorbance of cell lysates was measured at 570 nm. Error bars represent standard deviation from 10 wells.

### 3.3 Bioluminescence microscopy

The major limitation of the bacterial bioluminescence system for imaging applications is its comparatively low light emission. We have shown in previous work that the bioluminescence emission from mammalian cells expressing codon-optimized versions of the Lux proteins is sufficient for single-cell imaging [15]. However, exposure times of one to several minutes were required to obtain high-contrast images. Due to the limited photon emission rate, a trade-off between spatial and temporal resolution and image contrast has to be made. For instance, the bioluminescence signal can be collected in fewer larger pixels (resulting in decreased spatial resolution) if high imaging speed is required, or bioluminescence can be recorded for a longer period of time, allowing the observation of smaller cellular details at the expense of temporal resolution. For applications that involve both high spatial and temporal resolution, higher photon emission rates are thus required. To assess the benefit of the increase in light emission, we applied Akt2CA-P2A-RFK for bioluminescence microscopy of HeLa cells using different exposure times (Figure 6). As expected, the improved light emission from cells transfected with Akt2CA-P2A-RFK resulted in higher brightness and image contrast for a given exposure time. Hence, shorter exposure times are required with Akt2CA-P2A-RFK, which therefore allows image acquisition at increased speed while maintaining high spatial resolution and image contrast.

**Figure 6.**
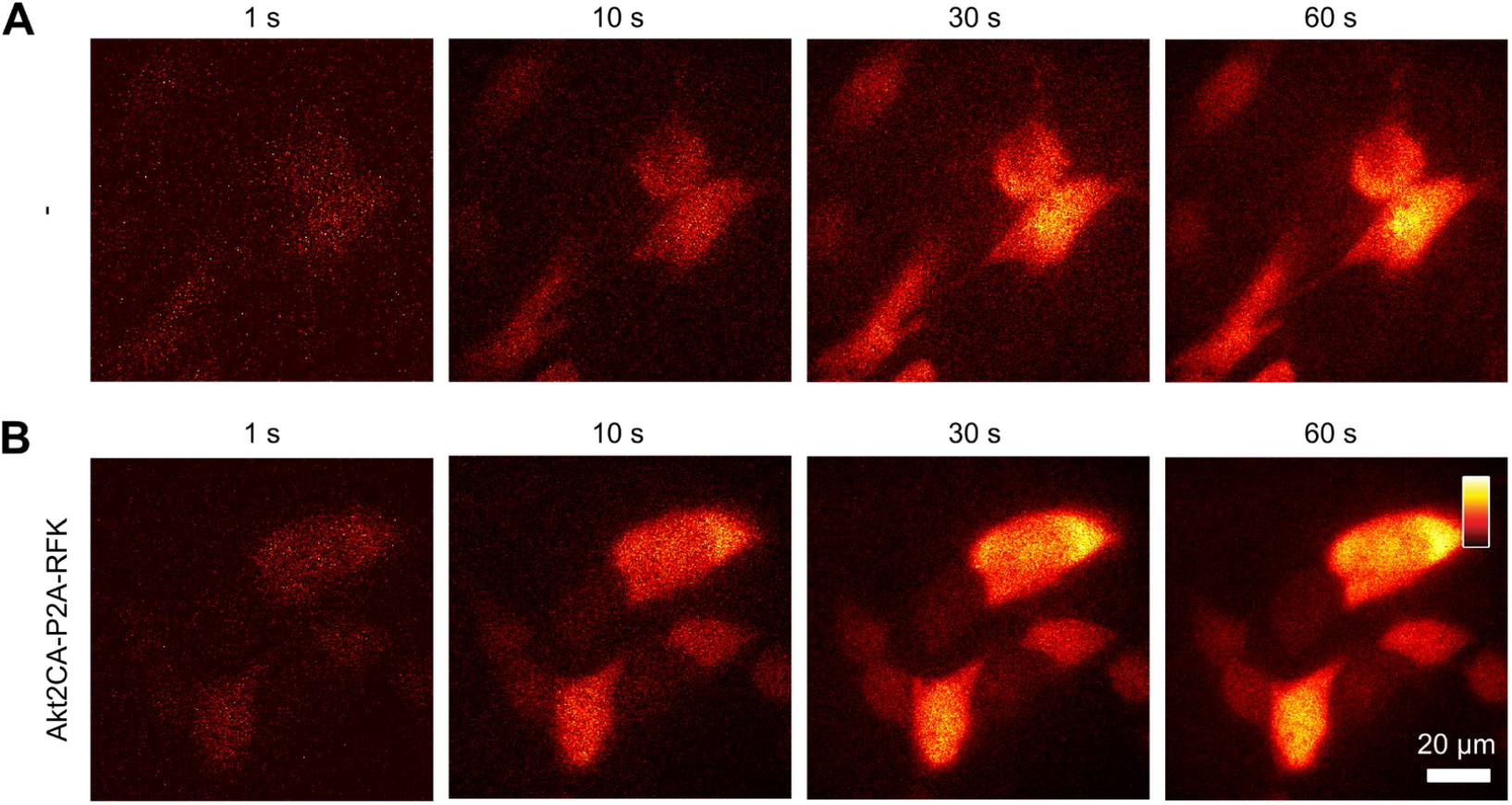
Bioluminescence of HeLa cells with and without Akt2CA-P2A-RFK. Cells grown on coverslips were transfected with a mixture of 0.8 µg lux plasmids and (**A**) 0.2 µg empty pcDNA3.1(+) vector or (**B**) 0.2 µg Akt2CA-P2A-RFK (all constructs in pcDNA3.1(+)). Bioluminescence emission was recorded using the indicated exposure times. The colormap was adjusted to the minimum and maximum pixel value of each image.

## 4. Discussion

The bacterial bioluminescence system is currently the only system that enables the generation of autonomous bioluminescence in mammalian cells. Problems associated with luciferin supply which is required for imaging using other bioluminescence systems are thereby circumvented. Besides, the light intensity of autonomous bioluminescence emission provides information on cell vitality since an active metabolism is required for continuous production of ATP and NADPH that are needed to recycle the substrates of the bioluminescence reaction. A major drawback of the bacterial bioluminescence system is its relatively low light output. This problem has been addressed previously by optimizing the expression of the Lux proteins [13–15]. In this way, increased light emission is achieved by higher cellular luciferase levels and enhanced production of the aldehyde and FMNH_2_ substrates from their direct precursors by LuxCDE and Frp, respectively.

Many applications of autonomous bioluminescence imaging would benefit from further increases in brightness. Based on the assumption that the concentrations of cofactors involved in the bioluminescence reaction may be limiting for light emission, we set out to increase the cellular levels of FMNH_2_ and NADPH. While FMNH_2_ is a direct substrate of the bacterial bioluminescence reaction, NADPH is required as a reductant for the regeneration of both the aldehyde and FMNH_2_ substrate. Compared to ATP that is also involved in substrate recycling, the concentration of NADPH in the cytosol of mammalian cells (3 µM [20]) is ∼3 orders of magnitude lower, making it more likely to be rate-limiting for the bioluminescence process. In bacteria, higher NADPH concentrations in the range of 100–300 µM have been reported [33– 35], and therefore this limitation may not occur in bioluminescent bacteria which represent the native cellular environment of the Lux proteins.

In this work, we therefore aimed to enhance FMNH_2_ and NADPH production in mammalian cell lines which increased bioluminescence levels up to 2.4–fold. The higher photon emission rate is beneficial for imaging applications where it improves image contrast and temporal and spatial resolution, as we demonstrated by bioluminescence microscopy of single cells. In addition, improved NADPH production by Akt2CA expression may be applied to increase light emission of other genetically encodable bioluminescence system such as the fungal system in the future which also require NADPH for substrate recycling.

## Author Contributions

Conceptualization, C.G.; methodology, T.B. and C.G.; validation, T.B. and C.G.; formal analysis, T.B. and C.G.; investigation, T.B. and C.G.; data curation, T.B. and C.G.; writing— original draft preparation, C.G.; writing—review and editing, T.B. and C.G.; visualization, T.B. and C.G.; supervision, C.G.; project administration, C.G.; funding acquisition, C.G. All authors have read and agreed to the published version of the manuscript.

## Data Availability Statement

Any data generated in this study are available from the corresponding authors upon reasonable request.

## Acknowledgments

pDONR223-RFK was a gift from William Hahn & David Root (Addgene plasmid # 23698; http://n2t.net/addgene:23698; RRID:Addgene_23698). G6PD/pRK5 was a gift from Xiaolu Yang (Addgene plasmid # 41521; http://n2t.net/addgene:41521; RRID:Addgene_41521). SIRT2 Flag was a gift from Eric Verdin (Addgene plasmid # 13813; http://n2t.net/addgene:13813; RRID:Addgene_13813). pDONR223-NADK was a gift from William Hahn & David Root (Addgene plasmid # 23747; http://n2t.net/addgene:23747; RRID:Addgene_23747). pDONR223-AKT1 was a gift from William Hahn & David Root (Addgene plasmid # 23752; http://n2t.net/addgene:23752; RRID:Addgene_23752). pLenti-FLAG-Akt2-CA was a gift from Weiping Han (Addgene plasmid # 64050; http://n2t.net/addgene:64050; RRID:Addgene_64050). iNap1 pcDNA3.1 Hygro(+) was a gift from Yi Yang (East China University of Science and Technology).

## References

1. Hall, M.P.; Unch, J.; Binkowski, B.F.; Valley, M.P.; Butler, L.B.; Wood, M.G.; Otto, P.; Zimmerman, K.; Vidugiris, G., Machleidt, T.; et al. Engineered luciferase reporter from a deep sea shrimp utilizing a novel imidazopyrazinone substrate. ACS Chem Biol 2012, 7, 1848–1857. doi: 10.1021/cb3002478

2. Lee, K.-H.; Byun, S.S.; Paik, J.-Y.; Lee, S.Y.; Song, S.H.; Choe, Y.S.; Kim, B.-T. Cell uptake and tissue distribution of radioiodine labelled D-luciferin: implications for luciferase based gene imaging. Nucl Med Commun 2003, 24, 1003–1009. doi: 10.1097/00006231-200309000-00009

3. Berger, F.; Paulmurugan, R.; Bhaumik, S.; Gambhir, S.S. Uptake kinetics and biodistribution of 14C-D-luciferin—a radiolabeled substrate for the firefly luciferase catalyzed bioluminescence reaction: impact on bioluminescence based reporter gene imaging. Eur J Med Mol Imaging 2008, 35, 2275–2285. doi: 10.1007/s00259-008-0870-6

4. Zhao, H.; Doyle, T.C.; Wong, R.J.; Cao, Y.; Stevenson, D.K.; Piwnica-Worms, D.; Contag, C.H. Characterization of coelenterazine analogs for measurements of Renilla luciferase activity in live cells and living animals. Mol Imaging 2004, 3, 43–54. doi: 10.1162/15353500200403181

5. Bhaumik, S.; Gambhir, S.S. Optical imaging of Renilla luciferase reporter gene expression in living mice. Proc Natl Acad Sci U S A 2002, 99, 377–382. doi: 10.1073/pnas.012611099

6. Inoue, Y.; Sheng, F.; Kiryu, S.; Watanabe, M.; Ratanakanit, H.; Izawa, K.; Tojo, A.; Ohtomo, K. Gaussia luciferase for bioluminescence tumor monitoring in comparison with firefly luciferase. Mol Imaging 2011, 10, 377–385. doi: 10.2310/7290.2010.00057

7. Cormier, M.J.; Kuwabara, S. Some observations on the properties of crystalline bacterial luciferase. Photochem Photobiol 1965, 4, 1217–1225. doi: 10.1111/j.1751-1097.1965.tb09308.x

8. Puget, K.; Michelson, A.M. Studies in bioluminescence. VII. Bacterial NADH:flavin mononucleotide oxidoreductase. Biochimie 1972, 54, 1197–1204. doi: 10.1016/s0300-9084(72)80024-5

9. Rodriguez, A.; Riendeau, D.; Meighen, E. Purification of the acyl coenzyme A reductase component from a complex responsible for the reduction of fatty acids in bioluminescent bacteria. Properties and acyltransferase activity. J Biol Chem 1983, 258, 5233–5237. doi: 10.1016/S0021-9258(18)32563-8

10. Rodriguez, A.; Wall, L.; Riendeau, D.; Meighen, E. Fatty acid acylation of proteins in bioluminescent bacteria. Biochemistry 1983, 22, 5604–5611. doi: 10.1021/bi00293a023

11. Carey, L.M.; Rodriguez, A.; Meighen, E. Generation of fatty acids by an acyl esterase in the bioluminescent system of Photobacterium phosphoreum. J Biol Chem 1984, 259, 10216–10221. doi: 10.1016/S0021-9258(18)90952-X

12. Kotlobay, A.A.; Sarkisyan, K.S.; Mokrushina, Y.A.; Marcet-Houben, M.; Serebrovskaya, E.O.; Markina, N.M.; Gonzalez Somermeyer, L.; Gorokhovatsky, A.Y.; Vvedensky, A.; Purtov, K.V.; et al. Genetically encodable bioluminescent system from fungi. Proc Natl Acad Sci U S A 2018, 115, 12728–12732. doi: 10.1073/pnas.1803615115

13. Close, D.M.; Patterson, S.S.; Ripp, S.; Baek, S.J.; Sanseverino, J.; Sayler, G.S. Autonomous bioluminescent expression of the bacterial luciferase gene cassette (lux) in a mammalian cell line. PLoS One 2010, 5, e12441. doi: 10.1371/journal.pone.0012441

14. Xu, T.; Ripp, S.; Sayler, G.S.; Close, D.M. Expression of a humanized viral 2A-mediated lux operon efficiently generates autonomous bioluminescence in human cells. PLoS One 2014, 9, e96347. doi: 10.1371/journal.pone.0096347

15. Gregor, C.; Pape, J.K.; Gwosch, K.C.; Gilat, T.; Sahl, S.J.; Hell, S.W. Autonomous bioluminescence imaging of single mammalian cells with the bacterial bioluminescence system. Proc Natl Acad Sci U S A 2019, 116, 26491–26496. doi: 10.1073/pnas.1913616116

16. Johannessen, C.M.; Boehm, J.S.; Kim, S.Y.; Thomas, S.R.; Wardwell, L.; Johnson, L.A.; Emery, C.M.; Stransky, N.; Cogdill, A.P.; Barretina, J.; et al. COT drives resistance to RAF inhibition through MAP kinase pathway reactivation. Nature 2010, 468, 968–972. doi: 10.1038/nature09627

17. Jiang, P.; Du, W.; Wang, X; Mancuso, A.; Gao, X.; Wu, M.; Yang, X. p53 regulates biosynthesis through direct inactivation of glucose-6-phosphate dehydrogenase. Nat Cell Biol 2011, 13, 310–316. doi: 10.1038/ncb2172

18. North, B.J.; Marshall, B.L.; Borra, M.T.; Denu, J.M.; Verdin, E. The human Sir2 ortholog, SIRT2, is an NAD+-dependent tubulin deacetylase. Mol Cell 2003, 11, 437–444. doi: 10.1016/s1097-2765(03)00038-8

19. Lim, C.Y.; Bi, X.; Wu, D; Kim, J.B.; Gunning, P.W.; Hong, W.; Han, W. Tropomodulin3 is a novel Akt2 effector regulating insulin-stimulated GLUT4 exocytosis through cortical actin remodeling. Nat Commun 2015, 6, 5951. doi: 10.1038/ncomms6951

20. tao, R; Zhao, Y.; Chu, H.; Wang, A.; Zhu, J.; Chen, X.; Zou, Y.; Shi, M.; Liu, R.; Su, N.; et al. Genetically encoded fluorescent sensors reveal dynamic regulation of NADPH metabolism. Nat Methods 2017, 14, 720–728. doi: 10.1038/nmeth.4306

21. Gregor, C.; Gwosch, K.C; Sahl, S.J.; Hell, S.W. Strongly enhanced bacterial bioluminescence with the ilux operon for single-cell imaging. Proc Natl Acad Sci U S A 2018, 115, 962–967. doi: 10.1073/pnas.1715946115

22. Yonezawa, A.; Masuda, S.; Katsura, T.; Inui, K.-I. Identification and functional characterization of a novel human and rat riboflavin transporter, RFT1. Am J Physiol Cell Physiol 2008, 295, C632–641. doi: 10.1152/ajpcell.00019.2008

23. Yamamoto, S.; Inoue, K.; Ohta, K.-Y.; Fukatsu, R.; Maeda, J.-Y.; Yoshida, Y.; Yuasa, H. Identification and functional characterization of rat riboflavin transporter 2. J Biochem 2009, 145, 437–443. doi: 10.1093/jb/mvn181

24. Yao, Y.; Yonezawa, A.; Yoshimatsu, H.; Masuda, S.; Katsura, T.; Inui, K.-I. Identification and comparative functional characterization of a new human riboflavin transporter hRFT3 expressed in the brain. J Nutr 2010, 140, 1220–1226. doi: 10.3945/jn.110.122911

25. Xiao, W.; Wang, R.-S.; Handy, D.E.; Loscalzo, J. NAD(H) and NADP(H) redox couples and cellular energy metabolism. Antioxid Redox Signal 2018, 28, 251–272. doi: 10.1089/ars.2017.7216

26. Wang, Y.-P.; Zhou, L.-S.; Zhao, Y.-Z.; Wang, S.-W.; Chen, L.-L.; Liu, L.-X.; Ling, Z.-Q.; Hu, F.-J.; Sun, Y.-P.; Zhang, J.-Y.; et al. Regulation of G6PD acetylation by SIRT2 and KAT9 modulates NADPH homeostasis and cell survival during oxidative stress. EMBO J 2014, 33, 1304–1320. doi: 10.1002/embj.201387224

27. Veech, R.L.; Eggleston, L.V.; Krebs, H.A. The redox state of free nicotinamide-adenine dinucleotide phosphate in the cytoplasm of rat liver. Biochem J 1969, 115, 609–619. doi: 10.1042/bj1150609a

28. Hedeskov, C.J.; Capito, K.; Thams, P. Cytosolic ratios of free [NADPH]/[NADP+] and [NADH]/[NAD+] in mouse pancreatic islets, and nutrient-induced insulin secretion. Biochem J 1987, 241, 161–167. doi: 10.1042/bj2410161

29. Pollak, N.; Niere, M.; Ziegler, M. NAD kinase levels control the NADPH concentration in human cells. J Biol Chem 2007, 282, 22562–22571. doi: 10.1074/jbc.M704442200

30. Oka, S.-I.; Titus, A.S.; Zablocki, D.; Sadoshima, J. Molecular properties and regulation of NAD+ kinase (NADK). Redox Biol 2023, 59, 102561. doi: 10.1016/j.redox.2022.102561

31. Hoxhaj, G.; Ben-Sahra, I.; Lockwood, S.E.; Timson, R.C.; Byles, V.; Henning, G.T.; Gao, P.; Selfors, L.M.; Asara, J.M.; Manning, B.D. Direct stimulation of NADP+ synthesis through Akt-mediated phosphorylation of NAD kinase. Science 2019, 363, 1088–1092. doi: 10.1126/science.aau3903

32. Berridge, M.V.; Tan, A.S. Characterization of the cellular reduction of 3-(4,5-dimethylthiazol-2-yl)-2,5-diphenyltetrazolium bromide (MTT): subcellular localization, substrate dependence, and involvement of mitochondrial electron transport in MTT reduction. Arch Biochem Biophys 1993, 303, 474–482. doi: 10.1006/abbi.1993.1311

33. Bennett, B.D.; Kimball, E.H.; Gao, M.; Osterhout, R.; Van Dien, S.J.; Rabinowitz, J.D. Absolute metabolite concentrations and implied enzyme active site occupancy in Escherichia coli. Nat Chem Biol 2009, 5, 593–599. doi: 10.1038/nchembio.186

34. Goldbeck, O.; Eck, A.W.; Seibold, G.M. Real time monitoring of NADPH concentrations in Corynebacterium glutamicum and Escherichia coli via the genetically encoded sensor mBFP. Front Microbiol 2018, 9, 2564. doi: 10.3389/fmicb.2018.02564

35. Shen, Y.-P.; Liao, Y.-L.; Lu, Q.; He, X.; Yan, Z.-B.; Liu, J.-Z. ATP and NADPH engineering of Escherichia coli to improve the production of 4-hydroxyphenylacetic acid using CRISPRi. Biotechnol Biofuels 2021, 14, 100. doi: 10.1186/s13068-021-01954-6

