## Supporting Information for "Increased autonomous bioluminescence emission from mammalian cells by enhanced cofactor synthesis"

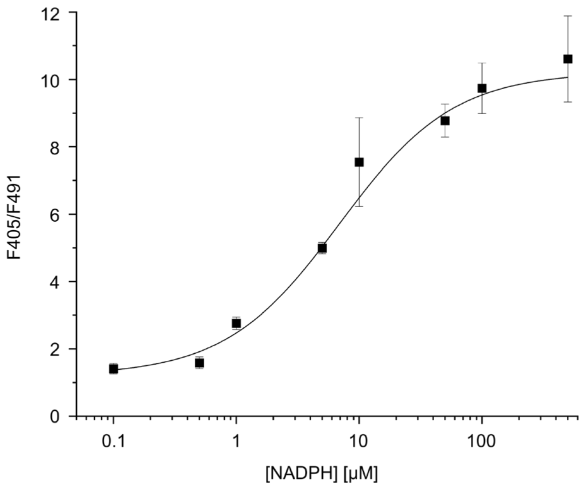


**Figure S1.** Calibration curve of iNap1. Fluorescence of purified iNap1 protein upon excitation at 405 (F405) and 491 nm (F491) was recorded at different NADPH concentrations. Data points and error bars represent average values and standard deviations of 5 mesurements. The measured values were fitted with a 4-parameter logistic (4PL) curve (shown as a line).

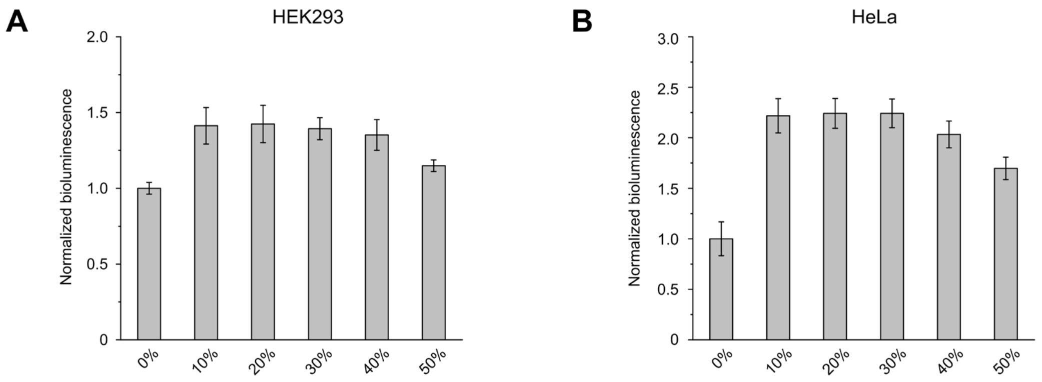


**Figure S2.** Brightness of cells transfected with lux plasmids and Akt2CA-P2A-RFK in different ratios. (**A**) HEK293 and (**B**) HeLa cells were grown in 24-well plates and transfected with a total amount of 0.5 µg DNA with the indicated percentage of the Akt2CA-P2A-RFK plasmid. Error bars represent standard deviation of 5 wells.

**Table S1.** Primers used for plasmid construction.

| Gene | Primer Name | Sequence (5'→3') |
|---|---|---|
| RFK | RFK NheI fwd | TGTAAAGCTAGCATGAGGCACCTGCCTTACTTC |
| | RFK XhoI rev | AATGTAGGCGCGCCTCAGTGGCCATTCATTATTTT |
| G6PD | G6PD NheI fwd | TGTAATGCTAGCATGGGTGCATCGGGTGAC |
| | G6PD XhoI rev | TGTAATCTCGAGTCAGAGCTTGTGGGGGTTCAC |
| SIRT2 | SIRT2 NheI fwd | AGTTTAGCTAGCATGGACTTCCTGCGGAAC |
| | SIRT2 XhoI rev | ATTGTACTCGAGTCACTGGGGTTTCTCCCT |
| NADK | NADK NheI fwd | ATGTAAGCTAGCATGGAAATGGAACAAGAAAAAATGA |
| | NADK BamH rev | GTAATAGGATCCCTAGCCCTCCTCCTCCTC |
| Akt1 | Akt1 NheI fwd | AGTAATGCTAGCATGAGCGACGTGGCTATT |
| | Akt1 XhoI rev | TGTAATCTCGAGTCAGGCCGTGCCGCTGGC |
| Akt2 | Akt2 NheI fwd | TGAATAGCTAGCATGAATGAGGTGTCTGTCAT |
| | Akt2 XhoI rev | TGAATACTCGAGTCACTCGCGGATGCTGGC |
| Akt1CA | myr-Akt1 NheI fwd | TGAATAGCTAGCATGGGTTCCTCCAAATCCAAGCCCAAGGCAAGCGCCATGATGAGCGACGTGGCTATTGTG |
| | Akt1 XhoI rev | TGTAATCTCGAGTCAGGCCGTGCCGCTGGC |
| Akt2CA | myr-Akt2 NheI fwd | TGAATAGCTAGCATGGGTTCCTCCAAATCCAAGCCCAAGGCAAGCGCCATGATGAATGAGGTGTCTGTCATCAAAGA |
| | Akt2 XhoI rev | TGAATACTCGAGTCACTCGCGGATGCTGGC |
| Akt2CA-P2A-RFK | myr-Akt2 NheI fwd | TGAATAGCTAGCATGGGTTCCTCCAAATCCAAGCCCAAGGCAAGCGCCATGATGAATGAGGTGTCTGTCATCAAAGA |
| | Akt2 BamHI rev | AATGTAGGATCCCTCGCGGATGCTGGCCGA |
| | P2A-RFK BamHI fwd | TTTTAGTGGATCCGGCGCCACCAACTTCAGCCTGCTGAAGCAGGCCGGCGACGTGGAGGAGAACCCCGGCCCCATGAGGCACCTGCCTTACTTC |
| | RFK XhoI rev | AATGTACTCGAGTCAGTGGCCATTCATTATTTT |
| RFK-P2A-Akt2CA | RFK NheI fwd | TGTAAAGCTAGCATGAGGCACCTGCCTTACTTC |

| | | |
|---|---|---|
| | RFK BamHI rev | AATGTAGGATCCGTGGCCATTCATTATTTTGCTTTTA |
| | P2A-myr-Akt2 BamHI fwd | TGAGTTGGATCCGGCGCCACCAACTTCAGCCTGCTGAAGCAGGCCGGCGACGTGGAGGAGAACCCCGGCCCCATGGGTTCCTCCAAATCCAAGCCC |
| | Akt2 XhoI rev | TGAATACTCGAGTCACTCGCGGATGCTGGC |
| His-iNap1 | His-iNap1 BamHI fwd | GTAATGGGATCCATGAGAGGATCGCATCACCATCACCATCACGGTTCTATGAACCGGAAGTGGGGC |
| | iNap1 SalI rev | AGTTACGTCGACCTAGCCCATCATCTCCTCCC |

**Table S2.** NADPH concentrations in Lux-expressing cells with and without Akt2CA expression. Cells were transfected and imaged as indicated in Figure 3. Cellular NADPH concentrations were calculated from the fluorescence ratio F405/F491 using the calibration curve shown in Figure S1.

| Cell line | Transfection | [NADPH] [μM] |
|---|---|---|
| LiveLight HEK293 | -Akt2CA | 9.3 |
| | +Akt2CA | 35.8 |
| HEK293 | -Akt2CA | 9.7 |
| | +Akt2CA | 14.0 |
| HeLa | -Akt2CA | 4.4 |
| | +Akt2CA | 11.8 |
